# Madariaga and Venezuelan equine encephalitis virus seroprevalence in rodent enzootic hosts in Eastern and Western Panama

**DOI:** 10.1101/2023.08.28.555226

**Authors:** Jean-Paul Carrera, Josefrancisco Galué, William M. de Souza, Rolando Torres-Cosme, Carlos Lezcano-Coba, Alberto Cumbrera, Nikos Vasilakis, Robert B. Tesh, Hilda Guzman, Scott C. Weaver, Amy Y. Vittor, Rafael Samudio, Juan Miguel Pascale, Anayansi Valderrama, Lorenzo Cáceres Carrera, Christl A. Donnelly, Nuno R. Faria

**Author notes:** Contributed equally (Jean-Paul Carrera, Josefrancisco Galué). These are joint senior authors in this study (Lorenzo Cáceres Carrera, Christl A. Donnelly, Nuno R. Faria).

## Abstract

While rodents are primary reservoirs of Venezuelan equine encephalitis virus (VEEV), their role in Madariaga virus (MADV) transmission remains uncertain, particularly given their overlapping geographic distribution. This study explores the interplay of alphavirus prevalence, rodent diversity, and land use within Darien and Western Panama provinces. A total of three locations were selected for rodent sampling in Darien province: Los Pavitos, El Real de Santa Maria and Santa Librada. Two sites were selected in Western Panama province: El Cacao and Cirí Grande. We used plaque reduction neutralization tests to assess MADV and VEEV seroprevalences in 599 rodents of 16 species across five study sites. MADV seroprevalence was observed at higher rates in Los Pavitos (Darien province), 9.0%, 95% CI: 3.6-17.6, while VEEV seroprevalence was elevated in El Cacao (Western Panama province), 27.3%, 95% CI: 16.1-40.9, and El Real de Santa María (Darien province), 20.4%, 95% CI: 12.6-29.7. Species like *Oryzomys coesi*, 23.1%, 95% CI: 5.0-53.8, and *Transandinomys bolivaris*, 20.0%, 95% CI: 0.5-71.6 displayed higher MADV seroprevalences than other species, whereas *Transandinomys bolivaris*, 80.0%, 95% CI: 28.3-99.4, and *Proechimys semispinosus*, 27.3%, 95% CI: 17.0-39.6, exhibited higher VEEV seroprevalences. Our findings provide support to the notion that rodents are vertebrate reservoirs of MADV and reveal spatial variations in alphavirus seropositivity among rodent species, with different provinces exhibiting distinct rates for MADV and VEEV. Moreover, specific rodent species are linked to unique seroprevalence patterns for these viruses, suggesting that rodent diversity and environmental conditions might play a significant role in shaping alphavirus distribution.

## Introduction

Madariaga (MADV) and Venezuelan equine encephalitic (VEEV) viruses (*Alphavirus* genus, *Togaviridae* family) are closely-related arthropod-borne zoonotic RNA viruses associated with the human and equine disease throughout Latin America ^1^. Most VEEV human-reported infections are symptomatic, and cases usually present with fever, headache, chills, and arthralgia ^2,3^. Around 14% of febrile cases develop severe neurological complications ^2^. VEEV case fatality ratio is estimated to be around 10% ^2^. MADV human infection is less well documented. In Panama, MADV was first reported in the former Panama Canal Zone in a horse in 1936 ^4^. Equine MADV epizootics were then reported across Panama, from the Azuero Peninsula in Central Panama to the Chepo district in North Panama, in 1947, 1958, 1962, 1973 and 1986 ^5–7^. An equine epizootic in the absence of human disease was also observed in Argentina in 1981 ^8^. In Iquitos, in the Peruvian Amazon, a febrile surveillance study found that 2% of participants were MADV IgM positive, indicating a low level of human exposure ^9^.

In 2010, 13 human MADV cases were reported during an outbreak of encephalitis in the Darien province, at the eastern end of Panama ^10^. Prior to this, a single case of human encephalitis had been reported in Brazil ^11^ and two MADV infections had been reported in Trinidad and Tobago ^12^. MADV human infections during the 2010 Panama outbreak presented with fever and headache, and rapidly developed neurological symptoms and complications ^10^ with an estimated case fatality ratio of around 10% ^10^. A recent report in Haiti showed that MADV human cases can present as a mild febrile disease with rash and conjunctivitis resembling symptoms observed during dengue disease ^13^. Similarly, human serosurveys undertaken in Panama suggested that the majority of MADV and VEEV infections are asymptomatic or cause mild disease ^3,14^. Nonetheless, follow-up studies of these individuals have demonstrated that clinical sequelae of MADV and VEEV can persist for years after infection^15^. Thus, the burden of both encephalitic alphaviruses could extend well beyond the acute febrile or neurological disease, such as described for arthritogenic alphavirus^16^. There are no VEEV- or MADV-specific treatments or licensed vaccines for use in humans. Diagnostic tests of human infections are typically performed using pan-alphavirus and/or virus-specific reverse transcription-polymerase chain reaction (RT-PCR) approaches, plaque reduction neutralization tests and viral isolation.

Mosquitoes within the subgenus *Culex (Melanoconion)* are believed to be the principal enzootic vectors of both VEEV and MADV. Previous studies in the Peruvian Amazon and Panama have shown frequent detection and isolation of MADV in *Culex* (*Mel*.) *pedroi* ^17,18^ *and Culex (Mel*.*) taeniopus taeniopus* ^7,19^. Furthermore, vector competence studies and analysis of blood feeding patterns show that *Culex (Mel*.) spp. predominantly feed on rodents in the wild ^2,18,19^. Indeed, experimental and field investigations suggest that several rodent species may act as host species for VEEV, including those within the genera *Sigmodon, Oryzomys, Zygodontomys, Heteromys, Peromyscus*, and *Proechimys* ^2,20,21^.

However, the vertebrate hosts for MADV remain poorly understood. Studies in wild rodents and marsupials in Brazil detected viremia in *Oryzomys* sp. (rice rat) and *Didelphis marsupialis* (common opossum) ^22–24^. MADV antibodies have also been detected in lizards and bats in Panama ^14,25^. Experimental studies in *Sigmodon hispidus* (cotton rat) and evolutionary analyses further support that rodent species may be a key amplifying host for MADV ^26,27^.

The geographic and temporal overlap of MADV and VEEV outbreaks in Panama suggests that these viruses occupy similar enzootic transmission cycles^10^. Recent studies suggest that rodent species collected in agricultural areas of Darien province were most likely to have MADV antibodies, while rodents with VEEV antibodies were principally found in sylvatic or forested areas ^14^. To elucidate the roles of distinct rodent species as hosts for alphaviruses, we conducted an assessment of MADV and VEEV seroprevalence within rodent populations. Additionally, we investigated the potential correlation between seroprevalence rates, rodent diversity, and the patterns of land use and land coverage across five distinct enzootic foci located in the Darien and Western Panama provinces.

## Materials and methods

### Ethics statement

The capture, use, and euthanization of wild rodents was evaluated and approved by the Institutional Animal Care and Use Committee of the Gorgas Memorial Institute for Health Studies (010/ CIUCAL/ICGES18) and the Panamanian Ministry of Environment (SC/A-21-17, ANAM) using the criteria established in the “International Guiding Principles for Biomedical Research Involving Animals” developed by the Council for International Organizations of Medical Sciences (CIOMIS). The study was conducted in accordance with Law No. 23 of January 15, 1997 (Animal Welfare Guarantee) of the Republic of Panama.

### Collection sites

Rodent trapping efforts were undertaken in 2011 and 2012 in Darien and Western Panama province (Figure 1). A total of three locations were selected for rodent sampling in Darien province: Los Pavitos, El Real de Santa Maria and Santa Librada (Figure 1). Two sites were selected in Western Panama province: El Cacao and Cirí Grande (Figure 1). The main economic activities in both regions are agriculture and cattle farming. Collection sites were selected based on previous reports of confirmed human and equine encephalitic alphavirus infection in 2001, 2004 and 2010^6^.

**Figure 1.**
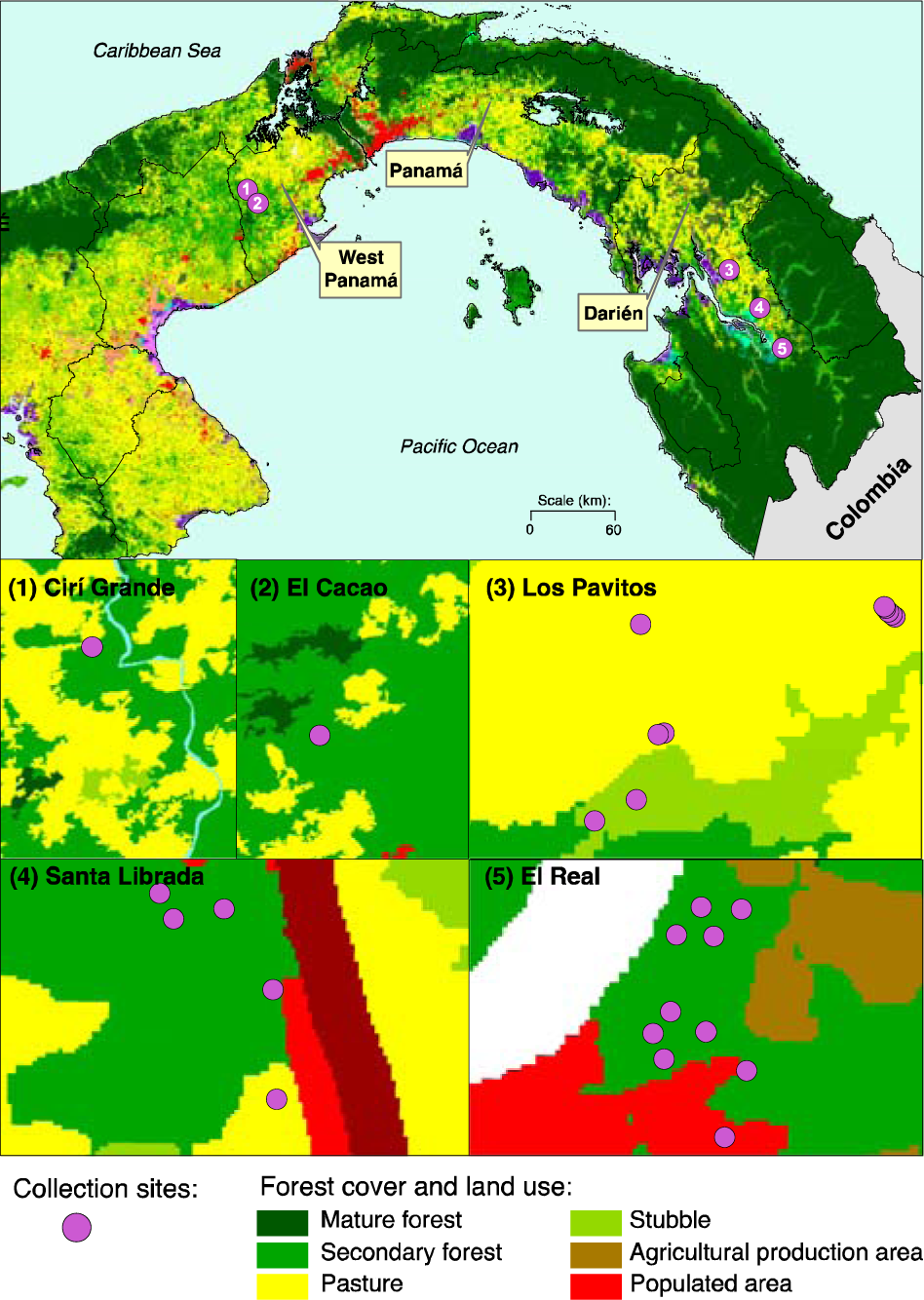
Study site and small mammal species. Study site map using the land use and land coverage (LULC) shapes ^46^. Classification of categories using the 2012 land use and land coverage shape. LULC categories were represented across all collection sites.

### Land use and land coverage classification

Georeferenced coordinates of collection sites were mapped onto the 2012 land use and land coverage (LULC) classification map obtained from the Panamanian Ministry of Environment ^28^ (Figure 1). The 2012 LULC classification was based on 5m resolution Rapid Eye Satellite Imagery ^29^.

### Small mammals trapping

From June to November 2011 and March to April 2012, small mammals were collected using Sherman traps baited with a mixture of rice, corn, sorghum, and peanut butter. In the field, traps were placed and maintained from 6:00 PM and then checked soon after 6:00 AM. For this study, a total of 100 Sherman traps were placed in three linear transects of approximately 125m during three consecutive nights at each location. Traps were placed within houses and in the peri-domiciliary area of previously confirmed VEEV cases. Peri-domiciliary setting includes grasslands, and crop fields as well as wooded areas near homes in each of the selected locations. Trapped animals were euthanized using halothane and identified using taxonomic keys or using the field guide to the mammals of Central America ^28^. Blood samples were collected from the retro-orbital sinus. Heart, liver, spleen, lung, and kidney tissues were then harvested. All samples were immediately placed into liquid nitrogen before transportation to the Gorgas Memorial Institute (GMI) for testing. Animal carcasses were deposited in the Vertebrate Museum of the University of Panama and the Zoological Collection of the GMI (Panama City, Republic of Panama).

### Laboratory methods

#### Alphavirus serology in small mammals

Rodent blood samples were screened in a 1:20 dilution using virus-specific plaque reduction neutralization tests (PRNTs) for VEEV and MADV viruses and then titred. A positive sample was considered as the reciprocal of the highest dilution that reduced plaque counts by >80% (plaque reduction neutralization test, PRNT_80_), as previously described ^14^. For PRNT, we used the wild-type MADV strain GML-267113, isolated from a fatal human case in Panama in 2017 ^30^, and the VEEV vaccine strain TC83. MADV and VEEV seroprevalence was estimated along with 95% confidence intervals (95% CIs) by mammalian species, year of collection, and collection site.

#### Viral isolation and molecular testing

Rodent tissues were used to prepare a 10% tissue suspension with 2 mL of minimum essential medium supplemented with penicillin and streptomycin, and 20% fetal bovine serum and homogenized using a Tissue Lyser (Qiagen, Hidden, Germany). After centrifugation at 17,709 x g for 10 minutes, 200 μL of the supernatant were inoculated into each of two 12.5 cm^2^ flasks of Vero cells (African green monkey-ATCC CCL-81, USA). Samples were passaged twice for cytopathic effect confirmation.

Rodent tissue and cell culture supernatant were used for viral RNA extraction using the QIaAmp RNA viral extraction kit (Qiagen, Valencia, CA) and tested for alphaviruses using reverse transcription-polymerase chain reaction (RT-PCR) assays, as previously described ^31^.

### Statistical methods

#### Diversity and similarity analysis

We estimated the absolute and relative abundance of small mammals in the collection sites of Darien and Western Panama provinces during 2011 and 2012. To compare the diversity of small mammals within collection sites we used the Shannon-Wiener index (H)^32^. Lower values of H correspond to lower diversity. We also used Simpson’s diversity index 1-D (SDI), which ranges from 0 (least diversity) to 1 (maximal diversity) ^33^. Margalef’s index was used to measure species richness, with higher values corresponding to greater species richness ^34^. Diversity analysis was undertaken using the statistical package PAST version 4.03^35^. Finally, a pairwise analysis of species by location was also undertaken. P-values and 95% CIs were adjusted for multiple comparisons using Tukey’s honestly significant difference (HSD) test, based on the possible pairs of means and studentized range distribution^36^.

#### Factors associated with alphavirus seroprevalence

Rodent species were grouped at the genus level to account for the small sample size. Rodent species, VEEV (n=296) and MADV (n=292) seropositivity, and LULC classification were used for univariate logistic regression analysis. To evaluate risk factors at the community and genus level, we conducted separate univariate analyses for MADV and VEEV; in each case, the outcome variable was the presence/absence of antibodies against the virus, as determined by a PRNT_80_ titer >1:20. The associations between each outcome and independent variable (community, genus and LULC) were estimated using logistic regression and were expressed as odds ratios (ORs). Univariable and multivariable ORs were calculated with 95% CIs. Statistical analyses were undertaken using the package STATA version 14.1 (College Station, TX).

## Results

### Rodent abundance across study sites

We collected a total of 559 rodents between 2011 and 2012, with specimens belonging to 13 genera and 16 species (Figure 2 A and B, Supplementary Table 1). Most rodents were captured during 2011 (71.8% of all collections, *n* = 430/599). In general, the majority of rodents were captured within the Darien Province (87.6% of all collections, *n* = 525/599), specifically in El Real (33.7%, *n* = 202/599), followed by Los Pavitos (27.6%, *n* = 165/599) and Santa Librada (26.4%, *n* = 158/599) (Supplementary Table 1).

**Figure 2.**
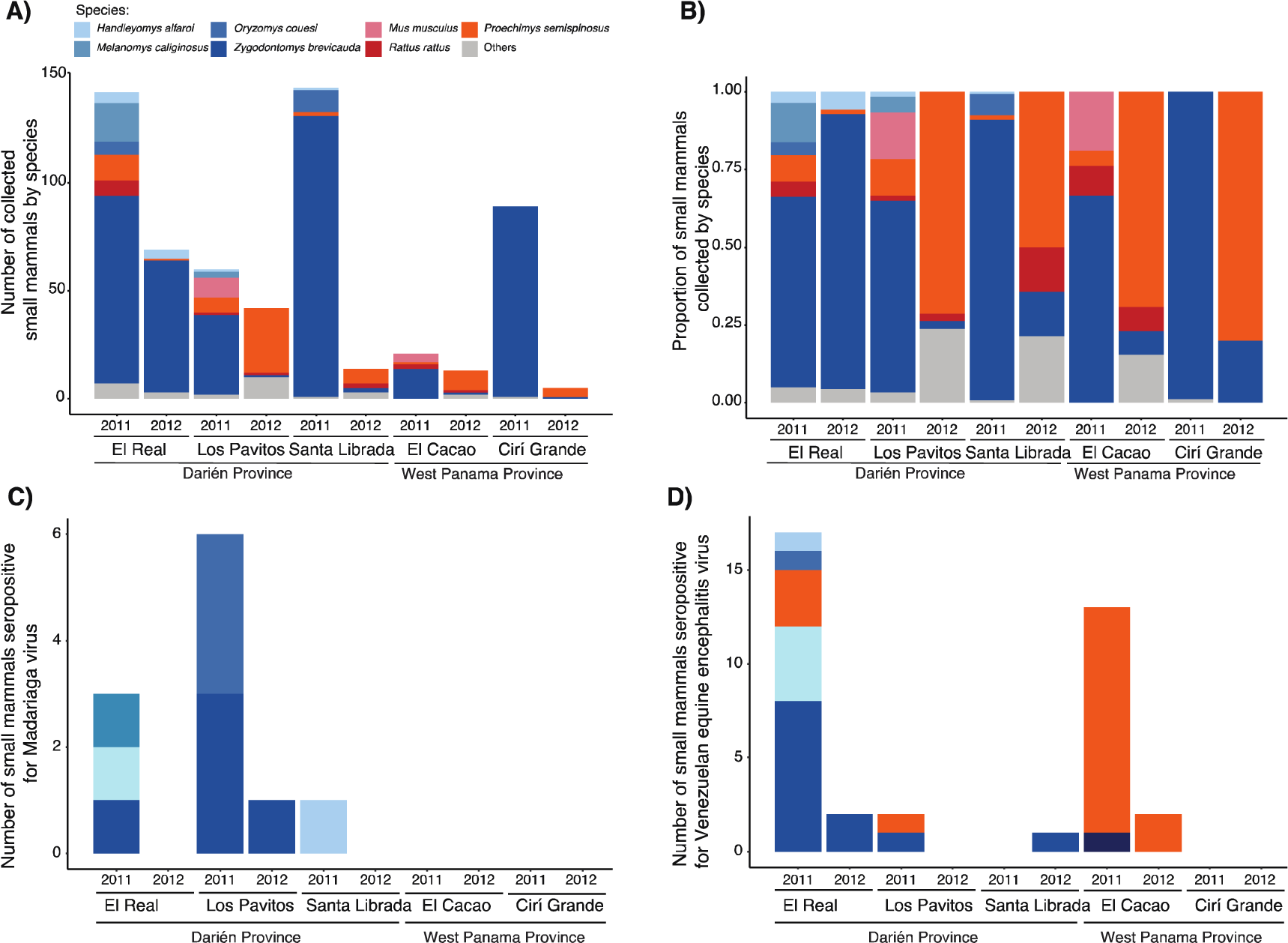
Alphavirus seropositivity in small mammals collected across study sites in Panama. A) Number of sampled small mammal species by site and year. B) Proportion of sampled small mammals by site and year. C) Number of small mammals seropositive for Madariaga virus (MADV). D) Number of small mammals seropositive for Venezuelan equine encephalitis virus (VEEV).

The short-tailed cane mouse (*Zygodontomys brevicauda*) was the most abundant species identified across study sites (70.5% of trapped animals, *n* = 402/599), followed by the Central American spiny rat (*Proechimys semispinosus*, 12.2%, *n =* 73/599), dusky rice rat (*Melanomys caliginosus*, 3.5%, *n* = 21/599), marsh rice rat (*Oryzomys couesi*, 2.7%, *n* = 16/599), the black rat (*Rattus rattus*, 2.3%), house mouse (*Mus musculus*, 2.2%, *n* = 13/599), Alfaro’s rice rat (*Handleyomys alfaroi*, 1.8%, *n* = 11/599), long-whiskered rice rat (*Transandinomys bolivaris*, 1.5%, *n* = 9/599), and the cotton rat (*Sigmodon hirsutus*, 1.3%, *n* = 8/599). Species with abundance ≤ 1% are shown in Supplementary Table 1.

### Highest rodent diversity and richness in the Darien Province

We estimated rodent diversity in each study site using the Simpson’s diversity index (1-D) and the Shannon-Wiener (H) index. The locations of El Real de Santa Maria [1-D=0.60; H=1.42] and El Cacao Maria [1-D=0.53; H=1.13] in the Darien province showed the highest rodent diversity. Lower species diversity was observed in Ciri Grande [1-D=0.46; H=0.96], Los Pavitos [1-D=0.23; H=0.57] and Santa Librada [1-D=0.11; H=0.29]. El Real de Santa Maria had the highest species richness accordingly with Margalef index [M=1.88] and Santa Librada presented the lowest species richness [M=0.79] (Table 1 and Supplementary Table 2).

**Table 1.**
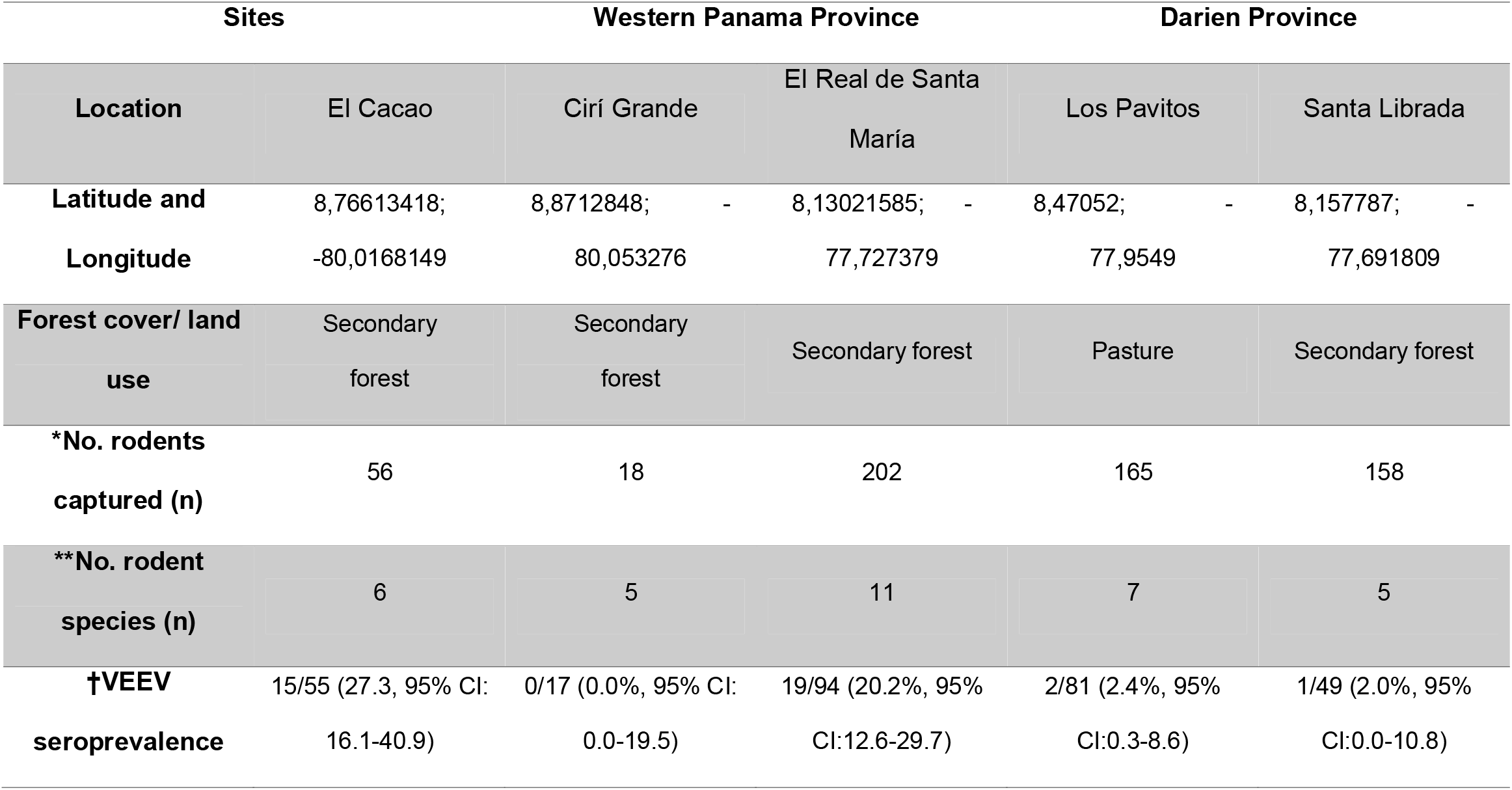
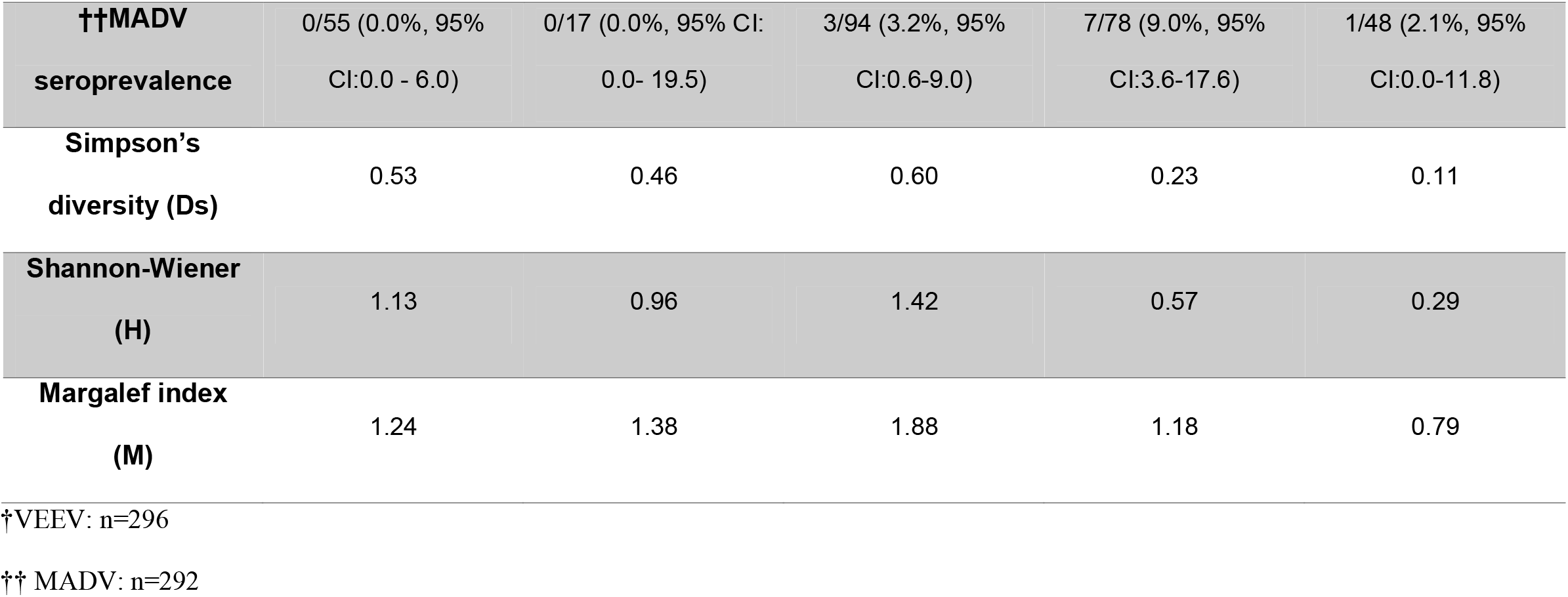
Characteristics of collection sites, small mammal diversity and alphavirus seroprevalence. The total of small mammals included in the analysis was 599 from a total of 16 species.

### Species similarity at the community level

Based on pairwise analyses, species composition was similar in Santa Librada and Los Pavitos in Darien province [Contrast =0.5; 95% CI: −0.5-1.4; p=0.639], and El Cacao and Ciri Grande in the Western province. Greater differences in species composition were observed between Darien and Western provinces (Table 3). Species compositions were generally most similar within provinces, with the exception of El Cacao and El Real de Santa Maria. These sites had the largest smallest differences in species composition [Contrast =-1.8; 95% CI: −3.1-0.5; p=0.001], despite these sites being in different provinces (Table 3).

**Table 2.**
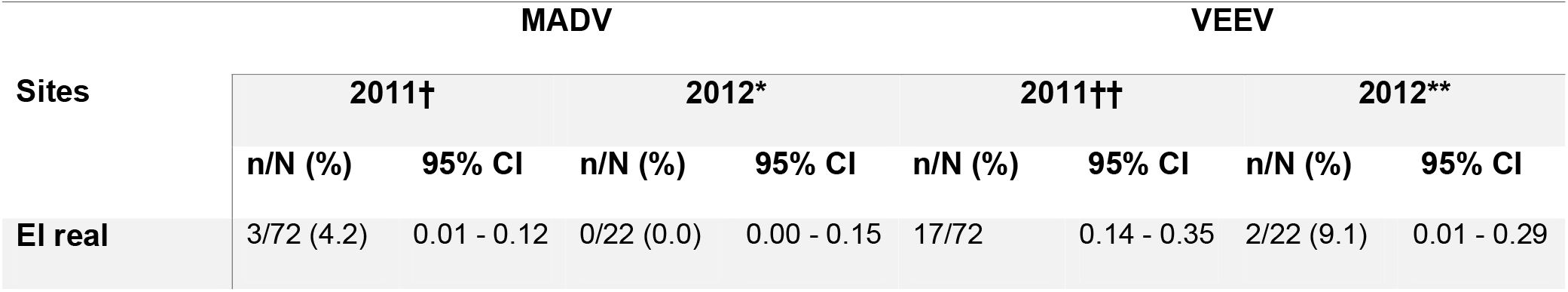
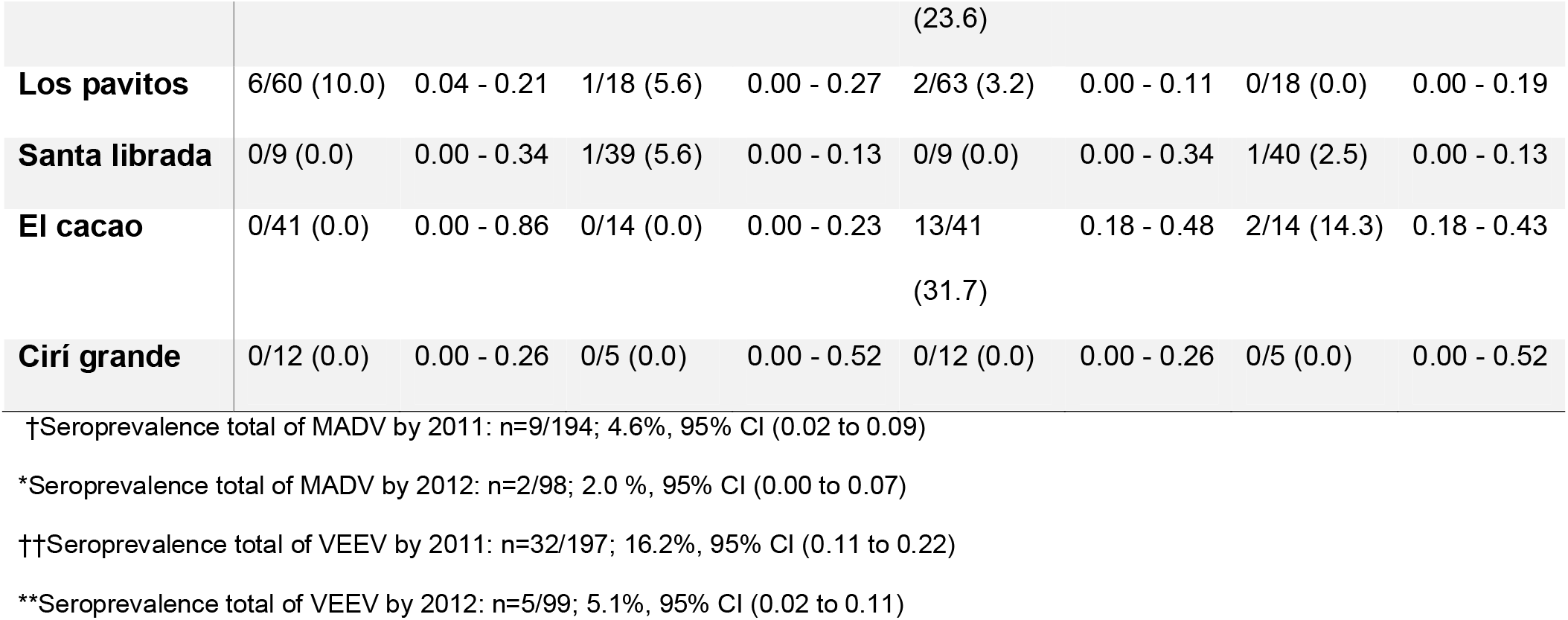
Seroprevalences by virus, collection sites and year of trapping.

**Table 3.** Pairwise comparison of rodent species by collection site.

| **Sites** | **Contrast** | **95% CI** | **P-value** |
| --- | --- | --- | --- |
| **Ciri Grande vs**<br>**El Real** | **-2.3** | **-4.4 - - 0.2** | **0.023** |
| **El Cacao vs El**<br>**Real** | **-1.8** | **-3.1- - 0.5** | **0.001** |
| **Los Pavitos vs**<br>**El Real** | **2.1** | **1.2 - 3.0** | **<0.001** |
| **Santa Librada**<br>**vs El Real** | **2.6** | **1.6 - 3.5** | **<0.001** |
| **El Cacao vs**<br>**Ciri Grande** | **0.5** | **-1.8 - 2.8** | **0.974** |
| **Los Pavitos vs**<br>**Ciri Grande** | **4.4** | **2.3 - 6.5** | **<0.001** |
| **Santa Librada**<br>**vs Ciri Grande** | **4.9** | **2.8 - 7.0** | **<0.001** |
| **Los Pavitos vs**<br>**El Cacao** | **3.9** | **2.6 - 5.2** | **<0.001** |
| **Santa Librada**<br>**vs El Cacao** | **4.4** | **3.0 - 5.7** | **<0.001** |
| **Santa Librada**<br>**vs Los Pavitos** | **0.5** | **-0.5 - 1.4** | **0.639** |

### Viral active circulation

No active alphavirus circulation was detected by means of RT-PCR or viral isolation. However, we note that two strains of Madrid virus (genus, *Orthobunyavirus*, family, *Peribunyaviridae*) were isolated from two specimens of *Zygodontomys brevicauda* trapped in El Real de Santa Maria. These strains are not analyzed in this study.

### Widespread alphavirus seroprevalence in rodents across Panama

The overall MADV and VEEV seroprevalence in small mammals were 3.8% (95% CI: 2.0-7.0; *n* = 11/292) and 12.5% (95% CI: 8.9-16.8; *n* = 37/296), respectively (Supplementary Table 3 and 4. VEEV seroprevalence was higher in 2011 (16.2%, 95% CI: 11.4-22.1; *n* = 32/197) compared to 2012 (5.1%, 95% CI: 1.6-11.3; *n* = 5/99) (Supplementary Table 6). MADV seroprevalence dropped from 4.6% (95% CI: 2.1-8.6; *n* = 9/194) in 2011 to 2.0% (95% CI: 0.2-7.0; *n* = 2/98) in 2012 (Supplementary Table 5). VEEV seroprevalence was widespread across the Western and Darien provinces with the highest seroprevalence found in El Cacao (27.3%, 95% CI: 16.1-40.9; n=15/55) in the Western province, followed by El Real de Santamaria (20.4%, 95% CI: 12.6-29.7; n=19/94) in the Darien province (Table 1, Table 2). MADV seroprevalence was higher in rodents collected in Los Pavitos (9.0%, 95% CI: 3.6-17.6 18; n=7/78), followed by El Real (3.2%, 95% CI: 1.0-9.0; n=3/94) and Santa Librada (2.1%, 95% CI: 0.0-11.0; n=1/48) (Table 1 and Table 2). No evidence of MADV viremia or antibodies was found in rodents collected in the Western province (0%, 95% CI: 0.0-5.0; n= 0/72).

**Table 4.** Univariable and multivariable logistic regression. Associated factors with MADV and VEEV seroprevalence.

| **Variables** | **MADV\*** |  |  | **VEEV\*** |  |  |
| --- | --- | --- | --- | --- | --- | --- |
| **Collection Site** | **OR** | **95% CI** | **P-value†** | **OR** | **95% CI** | **P-value†** |
| **El Real de Santa María** | **Ref.** | **-** | **-** | **Ref.** | **-** | **-** |
| **Los Pavitos** | **3** | **0.8 – 12.0** | **0.122** | **0.1** | **0.0 - 0.4** | **0.002** |
| **Santa Librada** | **0.6** | **0.1 - 6.4** | **0.708** | **0.1** | **0.0 - 0.6** | **0.017** |
| **El Cacao** | **-** | **-** | **-** | **1.5** | **0.7 - 3.2** | **0.323** |

| **Environment** |  |  |  |  |  |  |
| --- | --- | --- | --- | --- | --- | --- |
| **Secondary forest** | **Ref.** | **-** | **-** | **7.7** | **1.8 - 32.7** | **0.006** |
| **Pasture** | **5.2** | **1.5 - 18.2** | **0.01** | **Ref.** | **-** | **-** |
| **Genus** |  |  |  |  |  |  |
| ***Zygodontomys*** | **Ref.** | **-** | **-** | **Ref.** | **-** | **-** |
| ***Melanomys*** | **2.3** | **0.3 - 21.3** | **0.460** | **-** | **-** | **-** |
| ***Handleyomys*** | **5.0** | **0.5 - 49.7** | **0.170** | **2.0** | **0.2 - 18.4** | **0.524** |
| ***Transandinomys*\*\*** | 7.5 | 0.7 - 79.9 | 0.095 | 49.0 | 5.1 - 473.8 | 0.001 |
| ***Oryzomys*** | **9.0** | **1.9 - 43.2** | **0.006** | 1.0 | 0.1 - 8.5 | 0.985 |
| ***Proechimys*** | - | - | - | **4.6** | **2.1 - 10.2** | **<0.001** |
| ***Sigmodon*** | - | - | - | 1.8 | 0.2 - 15.4 | 0.614 |

MADV = Madariaga virus; OR = odds ratio; VEEV = Venezuelan equine encephalitis virus.
\*Based on plaque reduction neutralization test results.
\*\* Small sample size, n=5
†Results with $P < 0.05$ are shown in boldface type.

*Oryzomys couesi* (23.1%, 95% CI: 5.0-53.8; n=3/13) and *Transandinomys bolivaris* (20.0%, 95% CI: 0.5-71.6 72; n=1/5) had the highest MADV seroprevalence (Supplementary Table 3), while *Transandinomys bolivaris* (80.0%, 95% CI: 28.3-99.4; n=4/5) and *Proechimys semispinosus* (27.3%, 95% CI: 17.0-39.6; n=18/66) had the highest VEEV seroprevalence (Supplementary Table 4).

### Factors associated with alphavirus seroprevalence in rodents

MADV seroprevalence was independent of collection site, but Los Pavitos (OR=0.1; 95% CI: 0.0-0.4; p=0.002) and Santa Librada (OR=0.1; 95% CI: 0.0-0.6; p=0.017) were protective factors for VEEV seropositivity when compared with El Real de Santa María (Table 4). Univariate analysis by rodent taxa revealed that the odds of MADV seropositivity was 9.0 times greater in *Orizomys* (OR=9.0; 95%CI: 1.9-43.2; p=0.006) compared to the reference *Zygodontomys*. The odds of VEEV seropositivity in *Proechimys* (OR=4.6; 95%CI: 2.1-10.2; p<0.001) were significantly higher than in the reference, *Zygodontomys* (Table 4). At the univariate level, pasture was significantly associated with MADV seropositivity when compared to the secondary forest (OR=5.2; 95% CI: 1.5-18.2; p=0.01). In contrast, the risk of VEEV seropositivity was significantly decreased in pastures when compared with secondary forest (OR=0.1; 95% CI: 0.3 - 0.6; p=0.031) (Table 4).

## Discussion

Our findings support the hypothesis that wild rodents serve as reservoirs for both MADV and VEEV ^14,26,27^. Our results show that MADV seropositivity was confined to the Darien province, whereas VEEV seropositivity was pervasive across the examined study sites. Rodents captured within areas characterized by pasture exhibited an elevated likelihood of MADV seropositivity in contrast to those within secondary forest environments. Conversely, rodents captured within secondary forest areas displayed an increased likelihood of VEEV seropositivity. Overall, we observed that MADV seropositivity was lower in rodents compared to VEEV (3.8% vs. 12.5%). Our seroprevalence results agree with separate surveillance efforts carried out in other regions in the Darien province during 2012 ^14^. Higher VEEV seropositivity compared to MADV seropositivity in rodents has also been observed in mosquitoes and humans ^30,37^. Higher VEEV seroprevalence may be due to intrinsic differences among viral strains, variation in vector competence, viral competition within the vector, or asymmetric cross-protective immunity ^15,38^.

Weaver et al. has previously suggested that the genera with the greatest evidence of participation in the enzootic transmission of VEEV were *Sigmodon, Oryzomys, Zygodontomys* and *Proechimys* ^2,39^. We found that *Transandinomys bolivaris* and *Proechimys semispinosus* had the highest VEEV seroprevalence in Panama (80.0% and 27.3%, respectively). Both species have been implicated as VEEV reservoirs in prior studies ^2^. Moreover, the highest MADV seroprevalence was found in *Oryzomys couesi, Transandinomys bolivaris* and *Handleyomys alfaroi* (23.1%, 20.0% and 14.3%, respectively). We also observed that in different communities of the Darien province, *Zygodontomys brevicauda* and *Transandinomys bolivaris* presented the highest MADV seroprevalence (8.3% and 3.1%) ^14^.

*Proechimys semispinosus* and *Transandinomys bolivaris*, the rodent species identified in this study with the highest VEEV seroprevalence in the Darien province, are often found in secondary and primary forests ^14^. *Oryzomys couesi* and *Transandinomys bolivaris*, the rodents with the highest MADV seroprevalence, are found in grasslands and agricultural areas. *Oryzomys cousi* is a semi-aquatic species that is adaptable to different environmental conditions ^40,41^. Herbaceous habitats, permanent and semi-permanent wetlands appear to be an important factor for the distribution of this rodent ^40,41^. It is likely that this plasticity favors MADV transmission in pasture or agriculture settings. However, it is unclear if the ecological conditions found in Darien support the development of Culex (*Mel*.) spp., or possibly other bridge vectors. The ecological profiling of the *Cx*. (*Mel*.) *spp*., done during the 1970s, suggest these species develop their cycles in floating plant water^42^. More recent findings have discovered species near human settlements and in secondary forests^30,43^, suggesting changes in their ecology.

VEEV was more prevalent in rodents captured in the communities of El Cacao in the Panama Western province and in El Real de Santa Maria (27.3% and 20.2%) located in the Darien province. Rodent diversity and richness were also higher in El Real de Santa Maria and El Cacao. Notably, El Real de Santa María is also among the regions with the highest VEEV human incidence ^14,30^. Los Pavitos had the highest MADV rodent seroprevalence (9.0%), and we also observed that the risk of MADV increased in pasture compared with the secondary forest. Interestingly, Los Pavitos is a community on the Pan-American Highway where the first MADV human and equine cases were reported during the 2010 outbreak ^10^. Human serosurveys have shown that the risk of human VEEV infection is associated with activities in the forest, which supports a sylvatic cycle for VEEV ^14,30^. Previous studies have also shown that human MADV infection risk is associated with farming and cattle ranching activities, suggesting that MADV transmission occurs predominantly in areas with agricultural activity ^14,30^.

It is important to note that no MADV-seropositive rodents were observed in the El Cacao and Cirí Grande communities in the Western province of Panama. This observation is in agreement with recent serological evidence of MADV in rodents and humans being restricted to the Darien province ^10^. However, it is in contrast with pre-1990s reports of MADV showing widespread circulation across Panama ^5–7^. It is unclear why contemporary MADV transmission is limited to the Darién province, but perhaps these earlier outbreaks represented epizootic expansion from a stable enzootic focus in eastern Panama ^44^ Evidence of geographic expansion of MADV has also been previously observed in Panama^5,6^. High rates of MADV in rodents were recorded previously near El Real de Santa Maria in the small, heavily forested community of Pijibasal^14^. This community is in the Darien Gap National Park, suggesting that the MADV enzootic cycle also occurs in forested areas ^14^. Overall, MADV and VEEV seroprevalence levels appear to differ spatially, and our results suggest that MADV seroprevalence was greater in places with low rodent diversity and pasture, while VEEV seroprevalence was greater in places with rodent high diversity and secondary forest. However, cross-protection immunity has also been proposed as a potential mechanism to explain these differences ^14,15^

The limitations of this study include a lack of precise information on the environment where the rodents were collected, which means we could not describe the micro-ecological conditions linked to the distribution and prevalence of infection in rodents. Finer-scale analyses to understand the effect of land use and land cover in diversity and alphavirus seroprevalence are currently underway by our group using additional rodent data from Darien. Little volume of sample is also available for testing for alphaviruses in small animals, which makes laboratory testing challenging in some individuals or even other taxa. Moreover, future cross-sectional rodent surveys will allow us to identify the temporal drivers of transmission and improve our understanding of the seasonal dynamics of VEEV and MADV across Panama ^45^.

In summary, our study corroborates the hypothesis that wild rodents act as reservoirs for both MADV and VEEV, offering unique seropositivity patterns^14^. We observed distinct geographical distributions, with MADV seropositivity concentrated in the Darien province and VEEV seropositivity prevalent across the surveyed sites. *Transandinomys bolivaris* and *Proechimys semispinosus* exhibited the highest VEEV seroprevalence, while *Oryzomys couesi, Transandinomys bolivaris*, and *Handleyomys alfaroi* showcased elevated MADV seroprevalence. Furthermore, ecological differences in habitat preference were linked to seroprevalence patterns, with secondary forests associated VEEV with seropositivity and agricultural environments associated with MADV seropositivity.

Areas with lower rodent diversity and pasture environments correlated with increased MADV seropositivity. In contrast, regions characterized by higher rodent diversity and secondary forests were associated with heightened VEEV seroprevalence. These patterns align with observed human infection risks^14,30^, supporting the potential impact of rodent-driven transmission in specific ecological contexts.

## Acknowledgments

We wish to express appreciation to Yaneth Pittí, Isela Guerrero, David Beltran, and Julio Cisneros for technical support with laboratory testing. We also thank Fatima Rodriguez for funding administration. AV, JMP and LC are members of the Sistema Nacional de Investigation (SNI), SENACYT, Panama.

## Disclaimers

The opinions expressed by authors contributing to this journal do not necessarily reflect the opinions of the Gorgas Memorial Institute of Health Studies, The Panamanian Government, or the institutions with which the authors are affiliated.

Potential conflicts of interest. All Authors: No reported conflicts of interest. Conflicts that the editor consider relevant to the content have been disclosed.

## References

1. Navarro JC, Carrera JP, Liria J, Auguste AJ, Weaver SC., 2017. Alphaviruses in Latin America and the introduction of chikungunya virus. Human Virology in Latin America: From Biology to Control

2. Aguilar P v., Estrada-Franco JG, Navarro-Lopez R, Ferro C, Haddow AD, Weaver SC., 2011. Endemic Venezuelan equine encephalitis in the Americas: Hidden under the dengue umbrella

3. Carrera JP, Bagamian KH, Travassos Da Rosa AP, Wang E, Beltran D, Gundaker ND, Armien B, Arroyo G, Sosa N, Pascale JM, Valderrama A, Tesh RB, Vittor AY, Weaver SC., 2018. Human and equine infection with alphaviruses and flaviviruses in panamá during 2010: A cross-Sectional study of household contacts during an encephalitis outbreak. American Journal of Tropical Medicine and Hygiene

4. Kelser R.A., 1937. Equine encephalomyelitis in Panama. Veterinary bulletin : 19–21

5. Medina G, Gleiser CA, Mackenzei RB., 1965. Brote de encefalomielitis equina en la Republica de Panama. Boletín de la Oficina Sanitaria Panamericana 58: 390–394

6. Obaldía N, Dutary B, Clavel F, Zarate JL, Alvarez O, Evans E, Molina A, Serrano R, Villareal A, Boy RR, de Gracia A, Chalmers F, Vega B. J, George M, Saa E, et al., 1991. Encefalomielitis Equina del Este, Epizootia de 1986 en Panamá. Notas veterinarias 1: 4–7

7. Dietz WH, Galindo P, Johnson KM., 1980. Eastern equine encephalomyelitis in Panama: The epidemiology of the 1973 epizootic. American Journal of Tropical Medicine and Hygiene

8. Sabattini MS, Daffner JF, Monath TP, Bianchi TI, Cropp CB, Mitchell CJ, Aviles G., 1991. Localized eastern equine encephalitis in Santiago del Estero Province, Argentina, without human infection. Medicina (B Aires) 51

9. Aguilar P v., Robich RM, Turell MJ, O’Guinn ML, Klein TA, Huaman A, Guevara C, Rios Z, Tesh RB, Watts DM, Olson J, Weaver SC., 2007. Endemic eastern equine encephalitis in the Amazon region of Peru. American Journal of Tropical Medicine and Hygiene

10. Carrera J-P, Forrester N, Wang E, Vittor AY, Haddow AD, López-Vergès S, Abadía I, Castaño E, Sosa N, Báez C, Estripeaut D, Díaz Y, Beltrán D, Cisneros J, Cedeño HG, et al., 2013. Eastern Equine Encephalitis in Latin America. New England Journal of Medicine

11. Alice F., 1956. Infeccao humana pelo virus “leste” da encefalite equina. Bol Inst Biol da Bahia (Brazil) 3: 3–9

12. Corniou B, Ardoin P, Bartholomew C, Ince W, Massiah V., 1972. First isolation of a South American strain of Eastern Equine virus from a case of encephalitis in Trinidad. Trop Geogr Med 24

13. Lednicky JA, White SK, Mavian CN, el Badry MA, Telisma T, Salemi M, OKech BA, Beau De Rochars VM, Morris JG., 2019. Emergence of Madariaga virus as a cause of acute febrile illness in children, Haiti, 2015-2016. PLoS Negl Trop Dis

14. Vittor AY, Armien B, Gonzalez P, Carrera J-P, Dominguez C, Valderrama A, Glass GE, Beltran D, Cisneros J, Wang E, Castillo A, Moreno B, Weaver SC., 2016. Epidemiology of Emergent Madariaga Encephalitis in a Region with Endemic Venezuelan Equine Encephalitis: Initial Host Studies and Human Cross-Sectional Study in Darien, Panama. PLoS Negl Trop Dis 10

15. Carrera JP, Pittí Y, Molares-Martínez JC, Casal E, Pereyra-Elias R, Saenz L, Guerrero I, Galué J, Rodriguez-Alvarez F, Jackman C, Pascale JM, Armien B, Weaver SC, Donnelly CA, Vittor AY., 2020. Clinical and serological findings of madariaga and venezuelan equine encephalitis viral infections: A follow-up study 5 years after an outbreak in Panama. Open Forum Infect Dis

16. Doran C, Elsinga J, Fokkema A, Berenschot K, Gerstenbluth I, Duits A, Lourents N, Halabi Y, Burgerhof J, Bailey A, Tami A., 2022. Long-term Chikungunya sequelae and quality of life 2.5 years post-acute disease in a prospective cohort in Curaçao. PLoS Negl Trop Dis 16

17. Turell MJ, O’Guinn ML, Jones JW, Sardelis MR, Dohm DJ, Watts DM, Fernandez R, Travassos Da Rosa A, Guzman H, Tesh R, Rossi CA, Ludwig G v., Mangiafico JA, Kondig J, Wasieloski LP, et al., 2006. Isolation of Viruses from Mosquitoes (Diptera: Culicidae) Collected in the Amazon Basin Region of Peru. J Med Entomol 42: 891–898

18. Turell MJ, O’Guinn ML, Dohm D, Zyzak M, Watts D, Fernandez R, Calampa C, Klein TA, Jones JW., 2008. Susceptibility of peruvian mosquitoes to eastern equine encephalitis virus. J Med Entomol

19. Srihongse S, Galindo P., 1967. The isolation of eastern equine encephalitis virus from Culex (Melanoconion) taeniopus Dyar and Knab in Panama. Mosquito News 27: 74–76

20. Deardorff ER, Forrester NL, Travassos Da Rosa AP, Estrada-Franco JG, Navarro-Lopez R, Tesh RB, Weaver SC., 2009. Experimental infection of potential reservoir hosts with venezuelan equine encephalitis virus, Mexico. Emerg Infect Dis 15

21. Sotomayor-Bonilla J, Abella-Medrano CA, Chaves A, Álvarez-Mendizábal P, Rico-Chávez Ó, Ibáñez-Bernal S, Rostal MK, Ojeda-Flores R, Barbachano-Guerrero A, Gutiérrez-Espeleta G, Aguirre AA, Daszak P, Suzán G., 2017. Potential sympatric vectors and mammalian hosts of venezuelan equine encephalitis virus in Southern Mexico. J Wildl Dis 53

22. Lopes OS, Sacchetta LA., 1974. Epidemiological studies on Eastern Equine Encephalitis virus in Sao Paulo, Brazil. Rev Inst Med Trop Sao Paulo 16

23. Ferreira IB, Pereira LE, Rocco IM, Marti AT, de Souza LT, Iversson LB., 1994. Surveillance of arbovirus infections in the Atlantic Forest Region, State of São Paulo, Brazil. I. Detection of hemagglutination-inhibiting antibodies in wild birds between 1978 and 1990. Rev Inst Med Trop Sao Paulo 36

24. Grayson MA, Galindo P., 1969. Ecology of Venezuelan equine encephalitis virus in Panama. J Am Vet Med Assoc 155: 2141–5

25. Craighead JE, Shelokov A, Peralta PH., 1962. The lizard: A possible host for eastern equine encephalitis virus in panama. Am J Epidemiol

26. Arrigo NC, Paige Adams A, Watts DM, Newman PC, Weaver SC., 2010. Cotton rats and house sparrows as hosts for north and south american strains of eastern equine encephalitis virus. Emerg Infect Dis

27. Arrigo NC, Adams AP, Weaver SC., 2010. Evolutionary Patterns of Eastern Equine Encephalitis Virus in North versus South America Suggest Ecological Differences and Taxonomic Revision. J Virol

28. Anon., 2010. A field guide to the mammals of Central America & southeast Mexico. Choice Reviews Online

29. Sang H, Zhang J, Zhai L, Xie W, Sun X., 2015. Analysis of RapidEye imagery for agricultural land mapping. International Conference on Intelligent Earth Observing and Applications 2015

30. Carrera JP, Cucunuba ZM, Neira K, Lambert B, Pitti Y, Liscano J, Garzon JL, Beltran D, Collado-Mariscal L, Saenz L, Sosa N, Rodriguez-Guzman LD, Gonzalez P, Lezcano AG, Pereyra-Elias R, et al., 2020. Endemic and epidemic human alphavirus infections in eastern Panama: An analysis of population-based cross-sectional surveys

31. Sánchez-Seco MP, Rosario D, Quiroz E, Guzmán G, Tenorio A., 2001. A generic nested-RT-PCR followed by sequencing for detection and identification of members of the alphavirus genus. J Virol Methods

32. Ortiz-Burgos S., 2016. Shannon-weaver diversity index. Encyclopedia of Earth Sciences Series

33. Gregorius HR, Gillet EM., 2008. Generalized Simpson-diversity. Ecol Modell 211

34. Death R., 2021. Margalefs Index - Population Dynamics - Ecology Center

35. Hammer Ø, Harper DAT, Ryan PD., 2001. PAST: Paleontological statistics software package for education and data analysis. Palaeontologia electronica. Curr Sci 4

36. Nanda A, Mohapatra DrBB, Mahapatra APK, Mahapatra APK, Mahapatra APK., 2021. Multiple comparison test by Tukey’s honestly significant difference (HSD): Do the confident level control type I error. International Journal of Statistics and Applied Mathematics 6

37. Carrera J-P, Arauz D, Rojas A, Cardozo F, Stittleburg V, Galue J, Lezcano-Coba C, Vasilakis N, Pascale JM, Valderrama A, Donnelly CA, Faria NR, Waggoner JJ., 2022. Real-time RT-PCR for Venezuelan equine encephalitis complex, Madariaga and Eastern equine encephalitis viruses: application in clinical diagnostic and mosquito surveillance. medRxiv

38. Kantor AM, Lin J, Wang A, Thompson DC, Franz AWE., 2019. Infection Pattern of Mayaro Virus in Aedes aegypti (Diptera: Culicidae) and transmission potential of the virus in mixed infections with chikungunya virus. J Med Entomol 56

39. Weaver SC, Ferro C, Barrera R, Boshell J, Navarro JC., 2004. Venezuelan Equine Encephalitis

40. del Campo JTF, Olvera-Vargas M, Silla-Cortés F, Figueroa-Rangel BL, Iñiguez-Dávalos LI., 2022. Composition and structure of vegetation and tide regulate the occurrence of Oryzomys couesi and Hodomys alleni in mangrove forests of Laguna de Cuyutlán, West-Central Mexico. Wetl Ecol Manag 30

41. Eubanks BW, Hellgren EC, Nawrot JR, Bluett RD., 2011. Habitat associations of the marsh rice rat (Oryzomys palustris) in freshwater wetlands of southern Illinois. J Mammal 92

42. Galindo P, Adames AJ., 1973. Ecological Profile of Culex (Melanoconion) aikenii (Diptera: Culicidae), Vector of Endemic Venezuelan Encephalitis in Panama1. Environ Entomol 2

43. Torres R, Samudio R, Carrera JP, Young J, Maârquez R, Hurtado L, Weaver S, Chaves LF, Tesh R, Caâceres L., 2017. Enzootic mosquito vector species at equine encephalitis transmission foci in the República de Panama. PLoS One

44. Brault AC, Powers AM, Villarreal Chavez CL, Navarro Lopez R, Cachón MF, Gutierrez LFL, Kang W, Tesh RB, Shope RE, Weaver SC., 1999. Genetic and antigenic diversity among eastern equine encephalitis viruses from North, Central, and South America. American Journal of Tropical Medicine and Hygiene 61

45. Raghwani J, Faust CL, François S, Nguyen D, Marsh K, Raulo A, Hill SC, Parag K V., Simmonds P, Knowles SCL, Pybus OG., 2022. Seasonal dynamics of the wild rodent faecal virome. Mol Ecol

46. Solano M., 2012. Panama 2012 Forest Cover and Land Use

