## Supplementary material for "Madariaga and Venezuelan equine encephalitis virus seroprevalence in rodent enzootic hosts in Eastern and Western Panama": Supplmental Materials

**Ecological features of encephalitic alphavirus in Panama**

**Supplementary Table 1. Absolute and relative rodent abundance by year and collection site.**

| Species | Years‡ | | Collection sites† | | | | |
| --- | --- | --- | --- | --- | --- | --- | --- |
|  | **2011** | **2012** | **El Real** | **Los Pavitos** | **Santa Librada** | **El Cacao** | **Cirí Grande** |
|  | **n/N (%)** | **n/N (%)** | **n/N (%)** | **n/N (%)** | **n/N (%)** | **n/N (%)** | **n/N (%)** |
| *Didelphis marsupialis* | 1/430 (0.2) | 0/169 (0.0) | 0/202 (0.0) | 0/165 (0.0) | 0/158 (0.0) | 0/56 (0.0) | 1/18 (5.6) |
| *Liomys adspersus* | 4/430 (0.9) | 0/169 (0.0) | 0/202 (0.0) | 0/165 (0.0) | 0/158 (0.0) | 4/56 (7.1) | 0/18 (0.0) |
| *Marmosa robinsoni* | 1/430 (0.23) | 0/169 (0.0) | 1/202 (0.5) | 0/165 (0.0) | 0/158 (0.0) | 0/56 (0.0) | 0/18 (0.0) |
| *Marmosa sp.* | 1/430 (0.2) | 0/169 (0.0) | 0/202 (0.0) | 0/165 (0.0) | 0/158 (0.0) | 0/56 (0.0) | 1/18 (5.6) |
| *Melanomys caliginosus* | 18/430 (4.2) | 3/169 (1.8) | 21/202 (10.4) | 0/165 (0.0) | 0/158 (0.0) | 0/56 (0.0) | 0/18 (0.0) |
| *Mus musculus* | 0/430 (0.0) | 13/169 (7.7) | 9/202 (4.5) | 4/165 (2.4) | 0/158 (0.0) | 0/56 (0.0) | 0/18 (0.0) |
| *Handleyomys alfaroi* | 6/430 (1.4) | 5/169 (3.0) | 6/202 (3.0) | 1/165 (0.6) | 4/158 (2.5) | 0/56 (0.0) | 0/18 (0.0) |
| *Transandinomys bolivaris* | 6/430 (1.4) | 3/169 (1.8) | 5/202 (2.5) | 1/165 (0.6) | 3/158 (1.9) | 0/56 (0.0) | 0/18 (0.0) |
| *Oryzomys sp.* | 16/430 (3.7) | 0/169 (0.0) | 6/202 (3.0) | 10/165 (6.1) | 0/158 (0.0) | 0/56 (0.0) | 0/18 (0.0) |
| *Ototylomys phyllotis* | 0/430 (0.0) | 2/169 (1.2) | 2/202 (1.0) | 0/165 (0.0) | 0/158 (0.0) | 0/56 (0.0) | 0/18 (0.0) |
| *Proechimys semispinosus* | 53/430 (12.3) | 20/169 (11.8) | 19/202 (9.4) | 3/165 (1.8) | 1/158 (0.6) | 37/56 (66.1) | 13/18 (72.2) |
| *Rattus rattus* | 9/430 (2.1) | 5/169 (3.0) | 8/202 (4.0) | 2/165 (1.2) | 0/158 (0.0) | 3/56 (5.4) | 1/18 (5.6) |
| *Rattus sp.* | 2/430 (0.5) | 0/169 (0.0) | 1/202 (0.5) | 0/165 (0.0) | 1/158 (0.6) | 0/56 (0.0) | 0/18 (0.0) |
| *Sigmodon hirsutus* | 5/430 (1.2) | 3/169 (1.8) | 0/202 (0.0) | 0/165 (0.0) | 0/158 (0.0) | 8/56 (14.3) | 0/18 (0.0) |
| *Sigmodon sp.* | 1/430 (0.2) | 0/169 (0.0) | 0/202 (0.0) | 0/165 (0.0) | 0/158 (0.0) | 1/56 (1.8) | 0/18 (0.0) |
| *Zygodontomys brevicauda* | 307/430 (71.4) | 115/169 (68.1) | 124/202 (61.4) | 144/165 (87.3) | 149/158 (94.3) | 3/56 (5.4) | 2/18 (11.1) |
| *Total* | 430/599 (71.8) | 169/599 (28.2) | 202/599 (33.7) | 165/599 (27.5) | 158/599 (26.3) | 56/599 (9.3) | 18/599 (3.0) |
| ‡Rodent abundance total by year: 2011 n=430 and 2012 n=169; abundance total n=599 | | | |  |  |  |  |
| †Rodent abundance total by collation site: El Real n=202, Los Pavitos n=165, Santa Librada n=158, El Cacao n=56 and Cirí Grande n=18 | | | | | | | |

**Supplementary Table 2. Simpson, Shannon and Margalef diversity index.**

| Index | Collection sites | | | | | Environment | |
| --- | --- | --- | --- | --- | --- | --- | --- |
|  | **El Real** | **Los Pavitos** | **Santa Librada** | **El Cacao** | **Cirí Grande** | **Secondary forest** | **Pasture** |
| Simpson_1-D | **0.6** | **0.23** | **0.11** | **0.53** | **0.46** | **0.56** | **0.24** |
| Shannon_H | **1.42** | **0.57** | **0.29** | **1.13** | **0.96** | **1.37** | **0.59** |
| Margalef_M | **1.88** | **1.18** | **0.79** | **1.24** | **1.38** | **2.47** | **1.18** |

**Supplementary table 3. MADV seroprevalences by species and year of collection.**

| Species | MADV | | | | | | |
| --- | --- | --- | --- | --- | --- | --- | --- |
|  | **Total†** | | **2011*** | | | **2012**** | |
|  | **n/N (%)** | **95% CI** | **n/N (%)** | **95% CI** | | **n/N (%)** | **95% CI** |
| *Didelphis marsupialis* | **0/1 (0.0)** | **0.0 - 98.0** | **0/1 (0.0)** | **0.0 - 98.0** | | **0/0 (0.0)** | **0.0 - 100.0** |
| *Liomys adspersus* | **0/4 (0.0)** | **0.0 - 60.0** | **0/4 (0.0)** | **0.0 - 60.0** | | **0/0 (0.0)** | **0.0 - 100.0** |
| *Marmosa sp.* | **0/1 (0.0)** | **0.0 - 98.0** | **0/1 (0.0)** | **0.0 - 98.0** | | **0/0 (0.0)** | **0.0 - 100.0** |
| *Melanomys caliginosus* | **1/14 (7.1)** | **0.0 - 34.0** | **1/14 (7.1)** | **0.0 - 34.0** | | **0/0 (0.0)** | **0.0 - 100.0** |
| *Mus musculus* | **0/7 (0.0)** | **0.0 - 41.0** | **0/0 (0.0)** | **0.0 - 100.0** | | **0/7 (0.0)** | **0.0 - 41.0** |
| *Handleyomys alfaroi* | **1/7 (14.3)** | **0.0 - 58.0** | **0/3 (0.0)** | **0.0 - 71.0** | | **1/4 (25.0)** | **0.0 - 81.0** |
| *Transandinomys bolivaris* | **1/5 (2.0)** | **0.0 - 72.0** | **1/4 (25.0)** | **0.0 - 81.0** | | **0/1 (0.0)** | **0.0 - 98.0** |
| *Oryzomys couesi* | **3/13 (23.1)** | **0.0 - 54.0** | **3/13 (23.1)** | **0.0 - 54.0** | | **0/0 (0.0)** | **0.0 - 100.0** |
| *Proechimys semispinosus* | **0/66 (0.0)** | **0.0 - 5.0** | **0/50 (0.0)** | **0.0 - 6.0** | | **0/16 (0.0)** | **0.0 - 21.0** |
| *Rattus rattus* | **0/9 (0.0)** | **0.0 - 34.0** | **0/5 (0.0)** | **0.0 - 52.0** | | **0/4 (0.0)** | **0.0 - 60.0** |
| *Rattus sp.* | **0/2 (0.0)** | **0.0 - 84.0** | **0/2 (0.0)** | **0.0 – 84.0** | | **0/0 (0.0)** | **0.0 - 100.0** |
| *Sigmodon hirsutus* | **0/7 (0.0)** | **0.0 - 41.0** | **0/4 (0.0)** | **0.0 - 60.0** | | **0/3 (0.0)** | **0.0 - 71.0** |
| *Sigmodon sp.* | **0/1 (0.0)** | **0.0 - 98.0** | **0/1 (0.0)** | **0.0 - 98.0** | | **0/0 (0.0)** | **0.0 - 98.0** |
| *Zygodontomys brevicauda* | **5/155 (3.2)** | **1.0 - 7.0** | **4/92 (4.4)** | **1.0 - 11.0** | | **1/63 (1.6)** | **0.0 - 9.0** |
| †Seroprevalence of MADV, 3.8%, 95% CI: 2.0-7.0; *n* = 11/292  *Seroprevalence of 2011, 4.6%, 95% CI: 2.1-8.6; *n* = 9/194  **Seroprevalence of 2012, 2.0%, 95% CI: 0.2-7.0; *n* = 2/98 | | |  | |  |  |  |

**Supplementary table 4. VEEV seroprevalences by species and year.**

| Species | VEEV | | | | | | |
| --- | --- | --- | --- | --- | --- | --- | --- |
|  | **Total†** | | **2011*** | | | **2012**** | |
|  | **n/N (%)** | **95% CI** | **n/N (%)** | | **95% CI** | **n/N (%)** | **95% CI** |
| *Didelphis marsupialis* | **0/1 (0.0)** | **0.0 - 98.0** | **0/1 (0.0)** | | **0.0 - 98.0** | **0/0 (0.0)** | **0.0 - 100.0** |
| *Liomys adspersus* | **0/4 (0.0)** | **0.0 - 60.0** | **0/4 (0.0)** | | **0.0 - 60.0** | **0/0 (0.0)** | **0.0 - 100.0** |
| *Marmosa sp.* | **0/1 (0.0)** | **0.0 - 98.8** | **0/1 (0.0)** | | **0.0 - 98.0** | **0/0 (0.0)** | **0.0 - 100.0** |
| *Melanomys caliginosus* | **0/14 (0.0)** | **0.0 - 23.0** | **0/14 (0.0)** | | **0.0 - 23.0** | **0/0 (0.0)** | **0.0 - 100.0** |
| *Mus musculus* | **0/7 (0.0)** | **0.0 - 41.0** | **0/0 (0.0)** | | **0.0 - 100.0** | **0/7 (0.0)** | **0.0 - 41.0** |
| *Handleyomys alfaroi* | **1/7 (14.3)** | **0.0 - 58.0** | **1/3 (33.3)** | | **0.0 - 91.0** | **0/4 (0.0)** | **0.0 - 60.0** |
| *Transandinomys bolivaris* | **4/5 (80.0)** | **0.2 - 99.0** | **4/4 (100.0)** | | **0.4 - 100.0** | **0/1 (0.0)** | **0.0 - 98.0** |
| *Oryzomys sp.* | **1/13 (7.7)** | **0.0 - 36.0** | **1/13 (7.7)** | | **0.0 - 36.0** | **0/0 (0.0)** | **0.0 - 100.0** |
| *Proechimys semispinosus* | **18/66 (27.3)** | **0.1 - 40.0** | **16/50 (32.0)** | | **20.0 - 47.0** | **2/16 (12.5)** | **2.0 - 39.0** |
| *Rattus rattus* | **0/9 (0.0)** | **0.0 - 34.0** | **0/5 (0.0)** | | **0.0 - 52.0** | **0/4 (0.0)** | **0.00 - 60.0** |
| *Rattus sp.* | **0/2 (0.0)** | **0.0 - 84.0** | **0/2 (0.0)** | | **0.0 - 84.0** | **0/0 (0.0)** | **0.00 - 100.0** |
| *Sigmodon hirsutus* | **0/7 (0.0)** | **0.0 - 41.0** | **0/4 (0.0)** | | **0.0 - 60.0** | **0/3 (0.0)** | **0.00 - 71.0** |
| *Sigmodon sp.* | **1/1 (100.0)** | **3.0 - 100.0** | **1/1 (100.0)** | | **3.0 - 100.0** | **0/0 (0.0)** | **0.00 - 100.0** |
| *Zygodontomys brevicauda* | **12/159 (7.6)** | **4.0 - 13.0** | **9/95 (9.5)** | | **4.0 - 17.0** | **3/64 (4.7)** | **0.00 - 13.0** |
| †Seroprevalence, 12.5%, 95% CI: 8.9-16.8; n= 37/296)  *Seroprevalence of 2011, 16.2%, 95% CI: 11.4-22.1; n=32/197  **Seroprevalence of 2012, 5.1%, 95% CI: 1.6-11.3; *n* = 5/99 | | | |  |  |  |  |
